# Virulence is associated with daily rhythms in the within-host replication of the malaria parasite *Plasmodium chabaudi*

**DOI:** 10.1101/2023.08.22.554271

**Authors:** Alíz T. Y. Owolabi, Petra Schneider, Sarah E. Reece

## Abstract

Asexually replicating stages of most malaria (*Plasmodium* spp.) parasite species replicate synchronously within the red blood cells of their vertebrate host. Rhythmicity in this intraerythrocytic developmental cycle (IDC) enables parasites to maximise exploitation of the host and align transmission activities with the time of day that mosquito vectors blood feed. The IDC is also responsible for the major pathologies associated with malaria, and plasticity in the parasite’s rhythm can confer tolerance to antimalarial drugs. Both the severity of infection (virulence) and synchrony of the IDC vary across species and between genotypes of *Plasmodium*, yet this variation is poorly understood. Theory predicts that virulence and IDC synchrony are negatively correlated and we tested this hypothesis using two closely related genotypes of the rodent malaria model *Plasmodium chabaudi* that differ markedly in virulence. We also test the predictions that in response to perturbations to the timing (phase) of the IDC schedule relative to the phase of host rhythms (misalignment), the virulent parasite genotype recovers the correct phase relationship faster, incurs less fitness loss, and so, hosts benefit less from misalignment of the virulent genotype. Our predictions are partially supported; the virulent parasite genotype was less synchronous in some circumstances and recovered faster from misalignment. While hosts were less anaemic when infected by misaligned parasites, the extent of this benefit did not depend on parasite virulence. Overall, our results suggest that interventions to perturb the alignment between the IDC schedule and host rhythms, and increase synchrony between parasites within each IDC, could alleviate disease symptoms. However, virulent parasites, which are better at withstanding conventional antimalarial treatment, would also be intrinsically better able to tolerate such interventions.

## 1 Introduction

Circadian rhythms have evolved to allow organisms to anticipate environmental cycles that have 24-hour periodicity (Panda *et al*, 2002; Patke *et al*, 2020). The daily rhythms in behaviour and physiology exhibited by hosts impose a rhythmic environment on their parasites, and so can mediate host-parasite interactions (Carvalho Cabral *et al*, 2019; Westwood *et al*, 2019; Prior *et al*, 2020; Hunter *et al*, 2022). For example, the time of day when infection occurs can determine life-or-death outcomes of infection for taxonomically diverse hosts, including *Arabidopsis, Drosophila* and mice (Halberg *et al*, 1960; Stone *et al*, 2012; Hevia *et al*, 2015; Heipertz *et al*, 2018; Sengupta *et al*, 2019). Time of day effects of infection are in part explained by circadian clock control of the host’s immune responses; for example, in mammals, leukocyte abundance, phagocytic activity, and cytokine release all oscillate over the circadian cycle (Scheiermann *et al*, 2013). While such rhythms allow hosts to prepare effective defences against invading pathogens and parasites, host rhythms can also provide infectious agents with opportunities. For instance, invasion and replication of *Leishmania* parasites is facilitated by the accumulation of their host cells (macrophages) in the blood during the host’s rest phase (Kiessling *et al*, 2017). The filarial nematodes *Brugia malayi* and *Wuchereria bancrofti* sequester in tissues in a rhythmic manner, which may be to ensure they are only exposed to dangers of immune defences in the peripheral circulation at the time of day when their mosquito vectors are seek blood (Hawking, 1967; Pichon & Treuil, 2004). Parasites are not merely passive bystanders to environmental oscillations and consequently, some have evolved autonomy over their own rhythms. For example, the fungus *Botrytis cinerea*, uses a canonical transcription-translation feedback loop (TTFL) oscillator to schedule rhythms in factors used for host exploitation (Hevia *et al*, 2015). Rhythms in African trypanosomes and malaria (*Plasmodium*) parasites also fulfil some of the criteria for possessing endogenous oscillators but their molecular underpinnings remain mysterious (Rijo-Ferreira *et al*, 2017; Smith *et al*, 2020; Subudhi *et al*, 2020; Prior *et al*, 2021).

While rhythmicity in parasite activities is assumed to be advantageous, a population of parasites within a host all carrying out the same behaviour at the same time results in synchrony, which may be intrinsically costly (Greischar *et al*, 2014). For example, malaria parasites replicate asexually in the red blood cells (RBCs) of their mammalian hosts, in a process termed the intraerythrocytic developmental cycle (IDC). In every IDC, a small fraction of asexual stages commits to becoming sexual stages (gametocytes), which are responsible for transmission to a mosquito vector and, ultimately, onward transmission to a vertebrate host. The IDC in most malaria species involves parasites progressing synchronously and sequentially through three main developmental stages (rings, trophozoites, schizonts). In general, the role of rings is to remodel the RBC, trophozoites feed and grow, and schizonts are responsible for producing progeny (merozoites) which are released at the end of the IDC (schizogony) (Tuteja, 2007). Synchronous schizogony, which causes recurring fevers at 24-, 48- or 72-hour intervals, is a hallmark of many *Plasmodium* species (Hawking *et al*, 1968; Gautret *et al*, 1995; Simpson *et al*, 2002; O’Donnell *et al*, 2011; Mideo *et al*, 2013). Schizogony in the rodent malaria model *P. chabaudi* follows a 24-hour rhythm and is timed to align with host feeding-fasting rhythms (Hirako *et al*, 2018; Prior *et al*, 2018; Reece & Prior, 2018). Specifically, IDC completion occurs during the feeding window, i.e. the 12-hour dark period for nocturnal mouse hosts, and this temporal coordination between host and parasite is likely explained by when rhythmic host nutrients – including the food-derived essential amino acid isoleucine – peaks in the blood (Prior *et al*, 2021). Thus, the timing (phase) of the IDC is thought to facilitate resource acquisition, but parasites that are perfectly synchronised (i.e. a high amplitude rhythm) may inadvertently compete for access to nutrients. In addition to “crowding” causing competition for time-limited nutrients, highly synchronous bursting may cause merozoite progeny to hinder each other’s access to red blood cells (RBCs) (Greischar *et al*, 2014).

The costs of synchrony are likely to be greater when parasites are at high density and / or when hosts have reduced appetite due to infection symptoms. Malaria parasite genotypes vary in within-host replication rates and densities achieved in the blood, in manners related to their virulence (defined here as the degree of harm caused to the host). For malaria parasites, replication rate and virulence are related to immunogenicity (through host immunopathology), sequestration and cytoadherence to the microvasculature and clumping (rosetting) (Cockburn *et al*, 2004; Long & Graham, 2011; De Niz *et al*, 2016). Due to the higher densities of virulent genotypes, synchrony is more likely to cause inadvertent resource competition amongst closely genetically related parasites. Indeed, observational data (Simpson *et al*, 2002; Dobaño *et al*, 2007; Touré-Ndouo *et al*, 2009; Ciuffreda *et al*, 2020) and mathematical modelling (Greischar *et al*, 2014) predicts that the synchrony of the IDC is associated with virulence. For example, synchronous *P. falciparum* infections are sometimes associated with low parasitaemia (Ciuffreda *et al*, 2020), and the loss of synchrony during *P. chabaudi* infections coincides with high parasite density and RBC limitation (O’Donnell *et al*, 2022). These observations could be explained by synchrony limiting virulence. Put another way, avirulent parasites may garner the benefits of timing the IDC to align with host rhythms, with fewer costs of a trade-off imposed by synchrony.

We test whether the IDC rhythm is linked to virulence using two very closely related (isogenic) genotypes of the rodent malaria *P. chabaudi* for that differ substantially in virulence (Mackinnon & Read, 1999). Specifically, we hypothesise that the more virulent genotype, which depletes more host resources (including RBCs), should have a less tightly synchronised IDC. Following perturbation of the IDC schedule relative to the rhythm of the host (misalignment), parasites readily reschedule to regain alignment with host rhythms within 5-7 IDCs (Gautret *et al*, 1995; Subudhi *et al*, 2020; O’Donnell *et al*, 2022). Plasticity in the IDC rhythm involves shortening the duration of each IDC by approximately 2 hours (O’Donnell *et al*, 2022), likely achieved by downregulating the expression of Serpentine Receptor 10 (Subudhi *et al*, 2020). Virulent genotypes are better able to cope with adverse within-host conditions, including antimalarial drug treatment and competition in mixed-genotype infections (De Roode *et al*, 2005; Bell *et al*, 2006; Schneider *et al*, 2008, 2012). Thus, we also hypothesise that the more virulent genotype suffers less from being misaligned and realigns to the phase of host rhythms faster. Our predictions are partially supported; the more virulent genotype is less synchronous, does not experience an increase in synchrony during misalignment, and recovers the correct IDC rhythm faster. However, we do not find that the impact of misalignment on overall asexual or gametocyte densities depends on virulence, and while disease severity is reduced by misalignment this is not in a virulence-dependent manner. Given that the IDC is responsible for the severity of malaria infections and fuels the production of transmission stages, our data are in line with the suggestion that disrupting the timing but not synchrony (or increasing synchrony) of the IDC is a novel approach to treat infections and reduce transmission (Prior *et al*, 2020).

## 2 Methods

### Hosts and parasites

We used the synchronously replicating rodent malaria parasite species *Plasmodium chabaudi chabaudi*, which was originally isolated from the shining thicket rat *Grammomys poensis* (previously called *Thamnomys rutilans*) (Landau, 1965), though rodent malarias also use *Mus musculus* as a natural host (Boundenga *et al*, 2019). Experimental hosts were 8–13-week-old female C57BL/6Jcrl mice (bred in house), housed in a standard 12 hours light:12 hours dark photoschedule (lights on [Zeitgeber Time 0; ZT0] at 10 am GMT, lights off at 10 pm GMT [ZT12]) and given *ad libitum* access to food and drinking water (supplemented with 0.05% para-aminobenzoic acid to support parasite growth (Jacobs, 1964)). Donor mice were housed in either the standard or reversed (12 hours dark:12 hours light; lights on at 10 pm GMT [ZT0], lights off at 10 am GMT [ZT12]) photoschedules. All donor and experimental hosts entrained to their photoschedule for 3.5 weeks prior to infection. All procedures occurred in accordance with the UK Home Office regulations (Animals Scientific Procedures Act 1986; SI 2012/3039; project licence number 70/8546) and were approved by The University of Edinburgh.

Experimental hosts received 1x10^7^ red blood cells (RBCs) infected with ring stage parasites and diluted in citrate saline (0.85% w/v NaCl, 1.5% w/v trisodium citrate dihydrate), belonging to either the relatively avirulent parental clone CW839 (referred to as CW-0) or its virulent progeny clone CW840 (referred to as CW-VIR). Prior to our study, parasites from the CW-0 parent were serially passaged from the mice within each selection cycle that experienced the most severe weight loss, and after 11 cycles of selection, CW-VIR achieved a 2.61-fold higher peak parasitaemia and caused 1.23-fold lower mean RBC density and 1.68 fold more mean weight loss than CW-0 (Mackinnon & Read, 1999). Using avirulent and virulent parasites of the same genetic background in our experiment minimises the influence of other phenotypes that exhibit genetic variation and that could confound the influence of virulence on IDC characteristics (Pollitt *et al*, 2011; Birget *et al*, 2019).

### Experimental design

We designed a cross-factored experiment (Fig. 1) in which each genotype (CW-0 and CW-VIR) was introduced into experimental hosts whose rhythms followed the same timing as the IDC (‘aligned’) or had rhythms 12 hours out of synch to the IDC (‘misaligned’). Establishing aligned infections involved infecting experimental hosts at ZT0 with ring stage parasites from donors in the same photoschedule, whereas we established misaligned infections by infecting experimental hosts at their ZT12 with ring stage parasites from donors in the opposite (reversed) photoschedule to create an instantaneous 12-hour shift in the phase of host rhythms for parasites. This design also enabled aligned and misaligned parasites to be sampled at the same time points within their host’s circadian cycle.

**Fig. 1.**
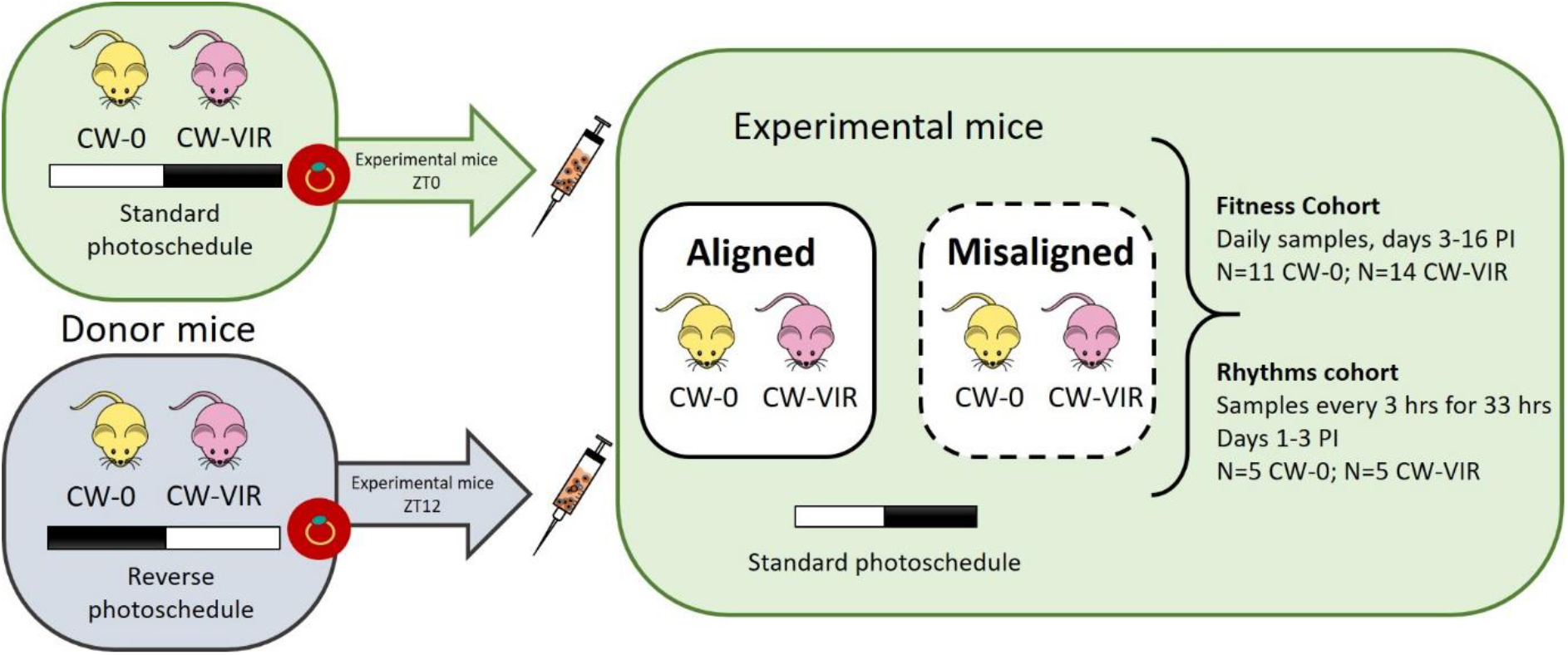
Experimental design. We housed donor mice in either standard (green, LD) or reversed (grey, DL) photoschedules, and all experimental mice were housed in the standard photoschedule. Mice received ring stage parasites of either the relatively avirulent *P. chabaudi* CW-0 (yellow) or the more virulent CW-VIR (purple) genotypes. To perturb the alignment of the intraerythrocytic developmental cycle (IDC) to host rhythms, we infected mice in ‘aligned’ groups at Zeitgeber Time (ZT) 0 (when lights went on) using parasites sourced from donors also at ZT0 in the standard photoschedule, and infected mice in the ‘misaligned’ groups at ZT12 (when lights went off), with parasites sourced from donors at ZT0 in the reverse photoschedule to create a 12-hour misalignment between the IDC schedule and host rhythms whilst standardising the IDC stage used to initiate infections. We replicated these four treatment groups in two cohorts, using the ‘Fitness cohort’ to assess parasite performance and disease severity, and the ‘Rhythms cohort’ to characterise the IDC schedule.

We initiated each of the four treatment groups in two cohorts of mice, termed the ‘Fitness’ and ‘Rhythms’ cohorts. Mice in the Fitness cohort (n = 11 per group for CW-0 infections and n = 14 per group for CW-VIR infections, Fig. 1) received their parasites by intraperitoneal injection and we followed their infections daily to measure the performance of parasites and disease severity. Mice in the Rhythms cohort (n = 5 per group for CW-0 infections and n = 6 per group for CW-VIR infections, Fig. 1) received their parasites by intravenous injection and we undertook intensive sampling within the first few IDCs to estimate parameters describing parasite rhythms. We used different cohorts for these purposes because the repeated sampling necessary to document IDC rhythms eventually generates anaemia and disrupts host rhythms, confounding assessment of parasite performance and virulence. Further, intravenous infection ensured parasites in the Rhythms cohort replicated to high enough densities to be quantifiable before those in misaligned infections had rescheduled, whereas intraperitoneal injection slows infection progression, avoiding the risk of mice in the Fitness cohort prematurely succumbing to infections. To initiate infections, blood from CW-0 or CW-VIR donors within each photoschedule was pooled to minimise donor effects and we randomly allocated experimental hosts to the four treatment groups and cohorts. We confirmed successful randomisation by measuring the weight and RBC density of each mouse the day before being infected (Suppl. Table 1).

### Sampling and data collection

We sampled the n = 50 infections in the Fitness cohort daily at ZT0 from day 3-16 post infection (PI) to quantify the densities of asexual stages and sexual transmission stages (gametocytes), and to track disease severity via weight and RBC density (Beckman-Coulter Counter). To determine total parasite density, we extracted DNA from 5 µl whole blood samples using a semi-automatic Kingfisher Flex Magnetic Particle Processor and MagMAX™-96 DNA Multi-Sample Kit (Thermo Fisher Scientific) with slight modifications from the standard protocol 4413021DWblood (Schneider *et al*, 2018a, 2019), followed by amplification of the CG2 gene (PCHAS_0620900, previously named PC302249.00.0) by qPCR (Wargo *et al*, 2007). To quantify gametocytes, we extracted RNA from 10 µl whole blood samples using the same Kingfisher machine and MagMAX™-96 Total RNA Isolation Kit (Birget *et al*, 2017; Schneider *et al*, 2019), followed by reverse transcriptase qPCR targeting the CG2 gene, which is expressed only in gametocytes (Wargo *et al*, 2006).

We sampled the n = 22 infections in the Rhythms cohort at 3-hourly intervals for 33 hrs from the ZT18 on the first day following infection (i.e. during 42-75 and 30-63 hours post infection (HPI) in the aligned and misaligned groups, respectively). The proportion of parasites at each IDC stage was quantified by staging 200 parasites per thin blood smear, stained with 20% Giemsa, based on their morphology following (Cambie *et al*, 1991; Prior *et al*, 2018). Technical issues precluded staging for the ZT12 sample, and where parasitaemia was very low (<0.2%; 29 out of 264 samples) at least 50 parasites were scored. Microscopic analysis was carried out blinded.

### Data analysis

To analyse parasite performance (total parasite and gametocyte densities) and disease severity (weight and RBC density), we compared dynamics over the course of infections as well as summary metrics comprising of peaks or minima, and cumulative counts. Only mice who survived to the end of the experiment, and thus, had a complete set of samples (n = 11 out of 11 for both groups of CW-0, and n = 10/14 for aligned and n = 13/14 for misaligned CW-VIR infections) were included in calculations of cumulative variables. The excluded mice reached the end point of infections on day 7 (n = 1), day 8 (n = 2) and day 12 (n = 1) PI in the aligned CW-VIR group, and day 8 PI (n = 1) in the misaligned CW-VIR group. All data from all mice were included in analyses of dynamics. For gametocyte density, we separately report densities for the first (day 3-8 PI) and the second peak (day 9-16 PI).

We characterised IDC rhythms from the proportion of asexual parasites at ring stage present in each sample from the Rhythms cohort (following (O’Donnell *et al*, 2011)) (Suppl. Fig. 1), using R (version 4.1.0) (R Foundation for Statistical Computing, Vienna, Austria, https://www.R-project.org/), and the packages ‘MetaCycle’ (function *meta2d*) (Wu *et al*, 2016) and ‘BiocManager’ (function *rain*) (Thaben & Westermark, 2014). The rhythmicity of ring stages was verified with both methods and only mice with *p* values < 0.05 (i.e. where the proportion of ring stages was deemed rhythmic by both functions) (n = 5 out if 5 per group for CW-0 and n = 4/6 per group for CW-VIR) were used to estimate period (the amount of time required to complete an IDC), phase (timing of the ring stage peak) and amplitude (the relative strength of the oscillation, as a proxy for synchrony of the IDC) for each infection individually using MetaCycle.

We used the R packages lme4 (for linear models) and lmer (for mixed effects models) to compare the effects of ‘Genotype’ (CW-0 or CW-VIR), ‘Alignment’ (aligned or misaligned to host rhythms) and their interaction. For analyses of dynamics, we also included ‘Day PI’ as a factor (to allow for nonlinear dynamics) as well as its 2- and 3-way interactions with genotype and alignment, and we account for pseudoreplication by including ‘Mouse ID’ as a random effect. We compared and minimised models using maximum likelihood deletion testing, prioritising lowest AICc (Akaike Information Criterion (corrected) for small sample sizes; ‘MuMIn’ package) when minimising non-nested parameters. Tests statistics and associated *p*-values are reported for significant results and full statistical results are shown in the Supplementary Tables. Effect sizes are given as fold differences (the quotients of the relevant means and 95% confidence intervals (CI) calculated using the Fieller method) or as mean ± standard error (SEM).

## 3 Results

### Verifying assumptions of the experimental design

In aligned infections, the two parasite genotypes *P. chabaudi* CW-0 and CW-VIR differed in virulence in the manner expected (Mackinnon & Read, 1999). CW-VIR reached higher peak parasite density, caused more severe anaemia and weight loss (Suppl. Fig. 2A-C), as well as higher mortality compared to CW-0. The perturbation of the alignment between parasite IDC schedule and host rhythm was successful for both genotypes, and misaligned parasites had not fully rescheduled their IDC to realign to their host’s rhythms during sampling of the “Rhythms cohort”. Specifically, the average proportion of ring stages for aligned infections peaks in the dark phase when the host feeds, but in the light phase for misaligned infections, reflecting the feeding phase of donor hosts (Suppl. Fig. 2D).

### Parasite performance

We expected that CW-0 parasites would experience a greater fitness cost of misalignment to host rhythms. To test this, we first compared parasite densities (as a proxy for within-host survival), finding that their dynamics differed between genotypes (χ^2^_(13)_ = 82.17, *p* < 0.001) and according to alignment (χ^2^_(13)_ = 278.50, *p* < 0.001; Suppl. Table 2A; Fig. 2A) but not their interaction. Specifically, CW-VIR exhibited higher densities, including exhibiting a second peak (day 12-13 PI), which CW-0 did not achieve. Across both genotypes, aligned parasites replicated to higher densities in the first few IDC than in misaligned infections (e.g. on day 4 PI, parasite density was 5.04-fold (95% CI 3.26, 7.69) higher in aligned infections; mean parasites per µl ± SEM = 9.79±1.48 ×10^4^ for aligned and 1.94±0.27 ×10^4^ for misaligned infections). However, across the whole infection, cumulative densities only reflect the difference between genotypes, with CW-VIR achieving 1.62-fold (95% CI 1.43, 1.84) higher densities (mean cumulative parasites per µl ± SEM = 7.98±0.18 ×10^5^ for CW-VIR and 4.94±0.29 ×10^5^ for CW-0) (*F*_(1,43)_ = 68.44, *p* < 0.001; Suppl. Table 2B; Fig. 2B).

**Fig. 2.**
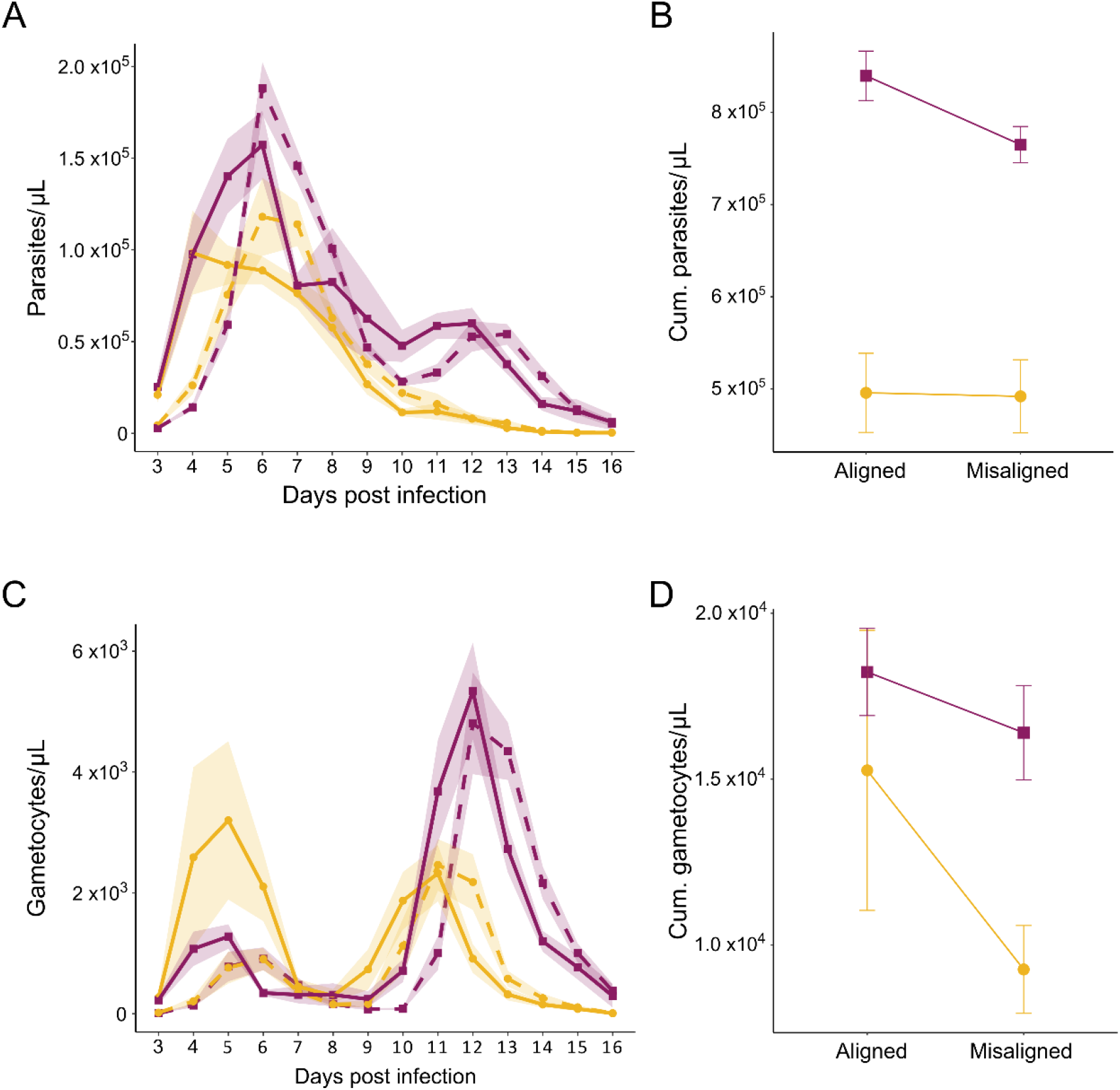
Parasite fitness proxies. The impact of alignment (solid lines) and misalignment (dashed lines) on CW-0 (yellow circles) and CW-VIR (purple squares) for measures of parasite performance (mean ± SEM): (**A**) total parasite density dynamics and (**B**) cumulative total parasite density, and (**C**) daily gametocyte dynamics and (**D**) cumulative gametocyte density. Cumulative densities are derived from summing data for individual hosts from day 3 to 16 PI.

Second, we compared gametocyte densities as a proxy for between-host transmission potential. The temporal dynamics of gametocyte densities differed between the genotypes in manners that depended on their alignment to host rhythms (χ^2^_(13)_ = 35.68, *p* = 0.001; Suppl. Table 2C; Fig. 2C). CW-VIR exhibited a more pronounced second peak in gametocyte densities, aligned infections followed patterns advanced by a day throughout infection, and misalignment only affected the magnitude of the early peak for CW-0. To further explore the impact of alignment and gametocyte on gametocyte dynamics, we separately analysed the maximum densities achieved in the first (days 3-8 PI) and second (days 9-16 PI) gametocyte peaks. Aligned infections produced 2.32-fold (95% CI 1.08, 4.02) higher gametocyte densities during the first peak compared to misaligned infections (mean peak gametocytes per µl ± SEM = 2.57±0.66×10^3^ for aligned and 1.11±0.17×10^3^ for misaligned infections) (*F*_(1,48)_ = 13.55, *p* < 0.001; Suppl. Table 2D; Fig. 2D), irrespective of genotype. However, only genotype explained the densities of the second gametocyte peak with CW-VIR achieving 1.94-fold (95% CI 1.48, 2.61) more gametocytes than CW-0 (mean peak gametocytes per µl ± SEM = 5.65±0.48×10^3^ for CW-VIR and 2.91±0.32×10^3^ for CW-0) (*F*_(1,43)_ = 22.55, *p* < 0.001; Suppl. Table 2E; Fig. 2D). Because the bulk of gametocytes were produced in the second peak, cumulative gametocyte densities reflected the difference between genotypes, with CW-VIR achieving 1.40-fold (95% CI 1.00, 2.25) higher densities (mean cumulative gametocytes per µl ± SEM = 1.72±0.10×10^4^ for CW-VIR and 1.23±0.23×10^4^ for CW-0) (*F*_(1,43)_ = 12.94, *p* < 0.001). In addition, we observe a borderline trend (*F*_(1,42)_ = 4.01, *p* = 0.052; Suppl. Table 2F; Fig. 2D) for misalignment reducing gametocyte density by 1.27-fold (95% CI 0.88, 1.74), with mean cumulative gametocytes per µl ± SEM = 1.67±0.23×10^4^ for aligned and 1.31±0.12×10^4^ for misaligned infections.

### Disease severity

We predicted that misalignment would reduce disease severity and that CW-0 parasites would be more affected by misalignment than CW-VIR and thus, CW-0 hosts would benefit disproportionately more from their parasites being misaligned. Host weight varied throughout infections, and these dynamics differed for hosts infected with each genotype (χ^2^_(13)_= 192.16, *p* < 0.001) and according to whether parasites were aligned or not with host rhythms (χ^2^_(13)_= 55.07, *p* < 0.001; Suppl. Table 3A; Fig. 3A) but not their interaction. As we observed for gametocytes and to some extent for total parasites, aligned infections followed dynamics advanced by 1 day compared to misaligned infections during the period of weight loss. Hosts infected with CW-VIR had 1.09-fold (95% CI 1.04, 1.15) lower minimum weights (mean minimum weight in grams ± SEM = 20.0±0.4 for CW-0 and 18.3±0.3 for CW-VIR), and regained their original weight later during recovery (Fig. 3A). Over the whole course of infection, any temporal effects of alignment resulted in no net impact on cumulative weight loss, yet the difference between genotypes remained with hosts of CW-0 retaining 1.05-fold (95% CI 1.01, 1.09) more weight than those infected with CW-VIR (mean cumulative weight in grams ± SEM =309.2±4.4 for CW-0 and 295.2±3.4 for CW-VIR) (*F*_(1,43)_ = 6.35, *p* = 0.016; Suppl. Table 3B; Fig. 3B).

**Fig. 3.**
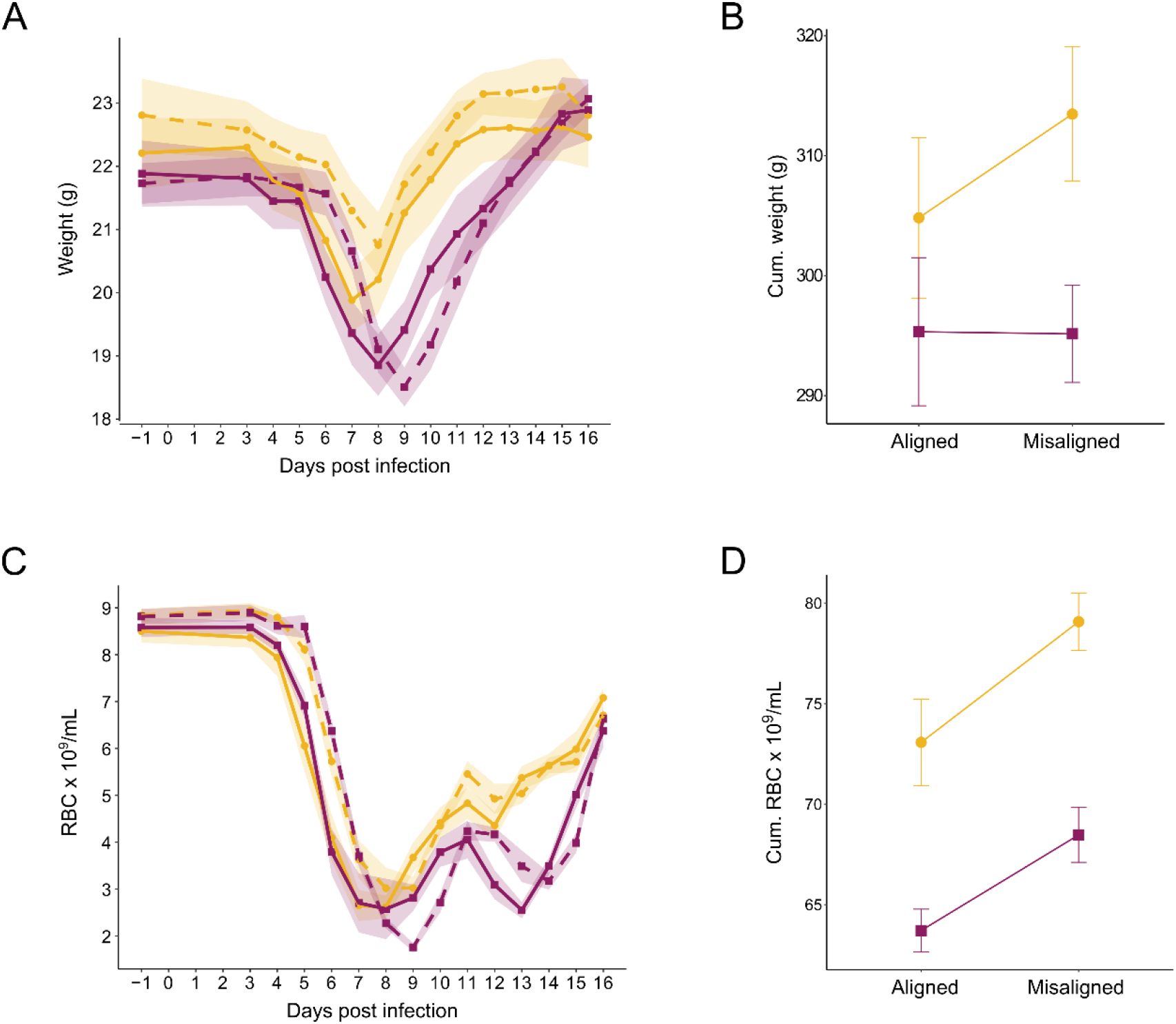
Disease severity proxies. Means ± SEM for aligned (solid lines) and misaligned (dashed lines) parasites of genotypes CW-0 (yellow circles) and CW-VIR (purple squares): (**A**) Mouse weights in grams from day -1 to 16 post infection (PI); (**B**) Cumulative weights, derived from summing data for individual hosts from day 3 to 16 PI; (**C**) Red blood cell (RBC) density per ml of blood from day -1 to 16 PI; and (**D**) Cumulative RBCs, derived from summing data for individual hosts from day 3 to 16 PI.

RBC dynamics also varied during infections in manners that depended on whether parasites were aligned or not (χ^2^_(13)_ = 120.68, *p* < 0.001) and on genotype (χ^2^_(13)_ = 122.84, *p* < 0.001), but not their interaction (Suppl. Table 3C; Fig. 3C). Mirroring the dynamics of most other variables, RBC densities of misaligned infections lagged behind those of aligned infections by a day and caused less cumulative anaemia, allowing their hosts to retain 1.07-fold (95% CI 1.005, 1.14) more RBCs over the course of infections (mean cumulative RBC per ml ± SEM = 73.33±1.47 ×10^9^ for misaligned and 68.62±1.60 ×10^9^ for aligned infections) (*F*_(1,42)_ = 11.94, *p* = 0.001; Suppl. Table 3D; Fig. 3D). Hosts infected with CW-VIR became more anaemic, with 1.38–fold (95% CI 1.16, 1.63) lower minimum RBC densities (mean minimum RBC per ml ± SEM = 2.36±0.13 ×10^9^ for CW-0 and 1.71±0.11 ×10^9^ for CW-VIR), experienced two pronounced troughs in RBC density (days 7-9 and 13-14 PI), regaining RBCs later (Fig. 3C), and retained 1.15-fold (95% CI 1.09, 1.20) less RBCs overall during infection (mean cumulative RBC per ml ± SEM = 76.08±1.42 ×10^9^ for CW-0 and 66.40±1.01 ×10^9^ for CW-VIR) (*F*_(1,42)_ = 41.78, *p* < 0.001; Suppl. Table 3D; Fig. 3D).

Finally, no CW-0-infected hosts reached the humane end point of infections, but in the CW-VIR infected hosts, 29% (n=4/14) with aligned parasites had to be euthanised compared to 7% (n=1/14) with misaligned infections.

### Characteristics of the IDC rhythm

We expected that CW-VIR parasites would be less synchronous than CW-0 in aligned infections and that CW-VIR would reschedule faster than CW-0 following misalignment. The oscillation of ring stages (Fig. 4A) was rhythmic in all CW-0 infections, but only in 8 of 12 CW-VIR infections (Suppl. Fig 1), suggesting CW-VIR is synchronous less often. Moreover, within the rhythmic infections, CW-VIR had a lower amplitude (i.e. was less synchronous) than CW-0 (mean amplitude = 0.43±0.03 for CW-0 and 0.22±0.02 for CW-VIR). This difference between genotypes was more pronounced in misaligned vs. aligned infections (*F*_(1,14)_= 38.64, *p* < 0.001; Suppl. Table 4A; Fig. 4B). Specifically, within aligned infections, the amplitude of ring stage proportions for CW-0 parasites was 1.38-fold (95% CI 1.01, 2.02) higher than for CW-VIR parasites (mean amplitude = 0.34±0.02 for aligned CW-0 and 0.25±0.03 for aligned CW-VIR), whilst this difference was 2.68-fold (95% CI 2.47, 2.92) in misaligned infections (mean amplitude = 0.53±0.01 for misaligned CW-0 and 0.20±0.01 for misaligned CW-VIR).

**Figure 4.**
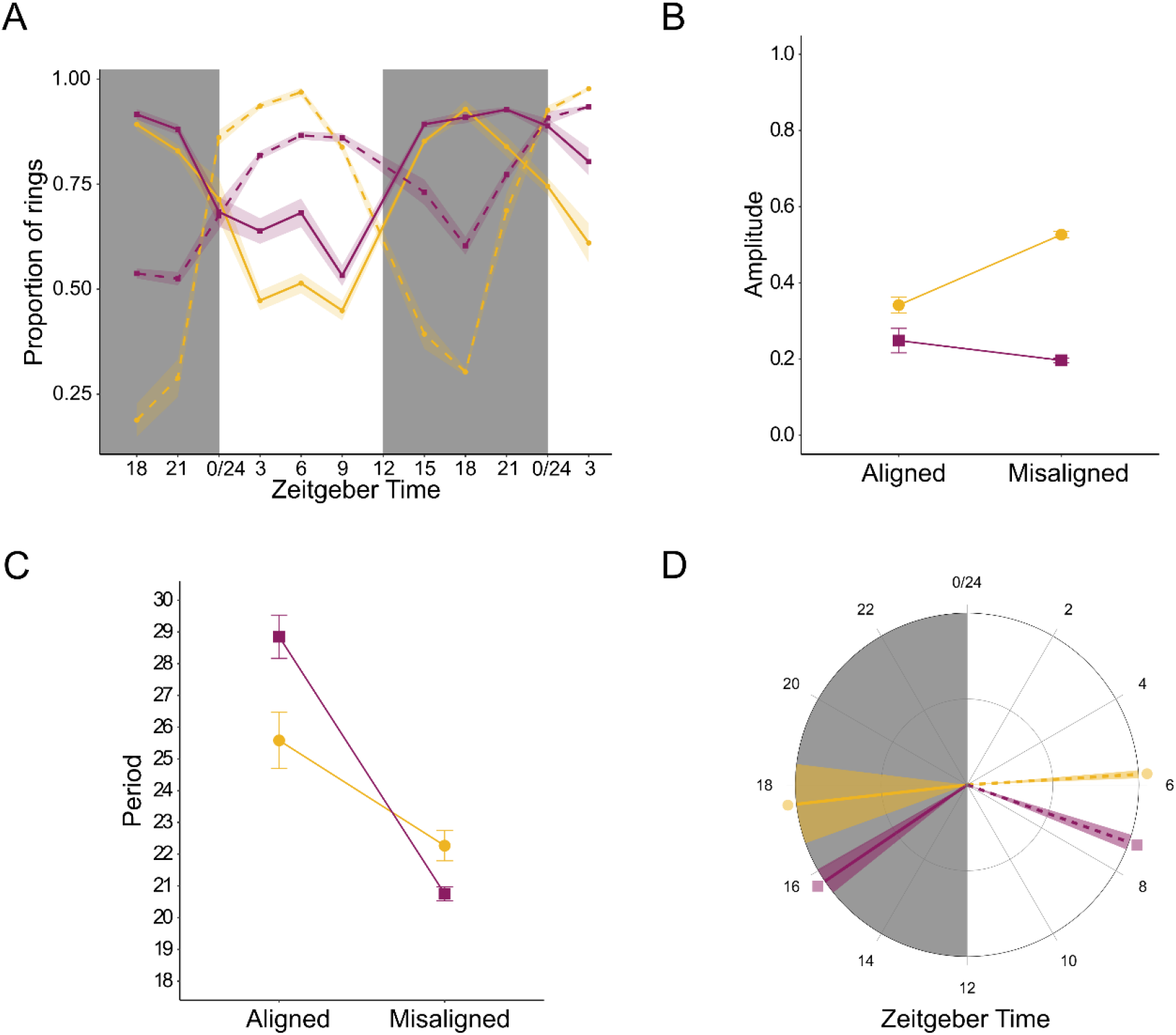
Intraerythrocytic developmental cycle (IDC) rhythms. Means ± SEM for aligned (solid lines) and misaligned (dashed lines) parasites of genotypes CW-0 (yellow circles) and CW-VIR (purple squares): (**A**) proportion of parasites at ring stage, (**B**) amplitude, (**C**) period, and (**D**) peak ring phase. Grey shading represents the dark phase of the circadian cycle (i.e. the host’s active phase).

When misaligned, parasites reschedule by shortening the IDC duration (Subudhi *et al*, 2020; O’Donnell *et al*, 2022), and this reduction was more pronounced for CW-VIR (mean difference = 8.1 hours, 95% CI 6.4, 9.8) than CW-0 (mean difference = 3.3 hours, 95% CI 1.0, 5.6) (*F*_(1,14)_ = 13.55, *p* = 0.002; Suppl. Table 4B; Fig. 4C). The ability of CW-VIR to reschedule its IDC more effectively is also reflected by the phase estimates for peak ring stages. Specifically, misaligned CW-VIR parasites were only 8.4 hours (95% CI 7.6, 9.3) out of phase to their aligned counterparts, but the phases for aligned and misaligned CW-0 were 11.8 hours (95% CI 9.7, 13.9) apart (*F*_(1,14)_ = 10.06, *p* = 0.007; Suppl. Table 4C; Fig. 4D).

## 4 Discussion

We tested whether virulence correlates with the synchrony of the IDC by comparing the impact of perturbing the IDC rhythm for two closely genetically related *P. chabaudi* genotypes, which vary in virulence: the relatively avirulent parental CW-0 and its descendent, the virulent CW-VIR (Mackinnon & Read, 1999). We predicted that CW-0 would suffer more than CW-VIR from the costs of being misaligned, resulting in greater loss of parasite and gametocyte densities and so, reducing the severity of infection symptoms. We confirmed that CW-VIR is more virulent than CW-0 and find some support for our predictions.

Misaligned infections exhibited approx. 1-day delayed dynamics for all parasite and host metrics (Fig. 2A, 2C, 3A, 3C). This may be explained – at least in part – by misaligned infections being initiated 12 hours after aligned infections. However, the significant interaction between alignment, genotype, and day PI on gametocyte dynamics coupled with lower early gametocyte peaks for both genotypes in misaligned infections suggests that in addition to any effect of infection age, misalignment has a direct negative impact on transmission potential (Fig. 2C). However, the second peak of gametocytes, which is not affected by alignment, ameliorates the impact of the early peak on overall transmission, especially for CW-VIR whose second gametocyte peak is much larger than its first. Misalignment reduced disease severity across infections by slowing weight loss (Fig. 3A) and curbing how much RBCs hosts lost (Fig. 3C-D), but these benefits were not greater for those infected with CW-0. Furthermore, CW-VIR was less synchronous, the amplitude of its IDC rhythm more robust to misalignment, and shortened its IDC duration more than CW-0 during rescheduling which brought CW-VIR’s phase closer to that of its aligned counterparts (Fig. 4). However, precise quantification, particularly of periods, from short time series data (as we collected), is challenging so we focus more on the qualitative than quantitative differences between genotypes for the characteristics of the IDC rhythm. Misalignment also reduced mortality from CW-VIR, but because no CW-0 infected mice reached their humane endpoints it is not clear whether misalignment causes a genotype specific difference in mortality. Overall, the impacts of misalignment are transient, becoming eroded as infections progress and reschedule to match the hosts’ rhythms (Fig. 2B, 2D). In contrast, the differences between the genotypes are consistent across parasite performance metrics and have greater impact than alignment.

Parasite fitness can be decomposed into traits that underpin within host survival and traits facilitating between-host transmission. Cycles of asexual replication maintain malaria in the host (and gametocytes usually make up less than 1% of parasites (Schneider & Reece, 2021)), thus, we use total parasite density as a proxy for within-host-survival and gametocyte density as an indicator of transmission potential (Cameron *et al*, 2013). Within-host survival and between-host transmission represent a resource allocation trade-off, because the production of asexual stages facilitates within-host survival via the accumulation parasite biomass but comes at a short-term cost of investment into gametocytes which are required for transmission to mosquitoes (Reece *et al*, 2009). Indeed, aligned and misaligned infections may illustrate this: by producing a lower early gametocyte peak (~57%, mean difference across genotypes = 1.46×10^3^ gametocytes per µl, 95% CI 0.10×10^3^, 2.83×10^3^), misaligned parasites’ replication is maximised, which could both ensure safety in numbers and provide a larger source population from which to invest in gametocytes later on in the infection (Westwood *et al*, 2020; Schneider & Reece, 2021). By reducing early gametocyte investment, parasites may ameliorate the costs of misalignment and achieve similar cumulative densities as their aligned counterparts (Fig. 2C, D) (Greischar *et al*, 2016; Schneider *et al*, 2018a). However, erosion of early transmission potential incurs a fitness cost when there is a risk of host mortality, even if modest (Greischar *et al*, 2016). Other non-mutually exclusive hypotheses for the negative impact of misalignment include elevated gametocyte mortality (O’Donnell *et al*, 2011) and sampling at the trough of the daily rhythm in gametocyte density due to their maturation and death rates (Pigeault *et al*, 2018; Schneider *et al*, 2018b). Future studies could combine high temporal resolution data with a within-host model of the maturation/senescence rates and density dynamics for gametocytes to explore the roles of plasticity in investment and mortality during misalignment and predict fitness outcomes.

Our readouts of disease severity are interlinked with parasite density dynamics; weight loss and anaemia become more severe as parasite densities increase, and mortality risk is highest following peak parasite density. Despite no clear impact of misalignment on cumulative total parasite density (Fig. 2B), hosts with misaligned parasites did experience less severe cumulative anaemia (Fig. 3D) and a lower mortality (for CW-VIR infections), and misalignment altered the temporal patterns of weight change (but not cumulative weight loss) (Fig. 3A-B). However, contrary to our expectation, hosts of avirulent parasites (CW-0) did not disproportionately benefit from the misalignment of their parasites. This may be due to misaligned CW-0 parasites prioritising asexual replication (via reducing investment into early gametocytes), so they do not disproportionately suffer from misalignment. The relatively minor impacts of misalignment exhibited by both genotypes suggests that early infection processes, potentially including time of day of infection (Stone *et al*, 2012; O’Donnell *et al*, 2014; Hevia *et al*, 2015; Sengupta *et al*, 2019), have long-lasting effects for RBC dynamics. For example, more RBCs are lost due to immunopathology than malaria parasite replication, and immunogenicity might be altered when infections occur at different times in the host’s circadian cycle, (Carvalho Cabral *et al*, 2022), and by undertaking cytoadherence and sequestration behaviours at an unusual time of day (i.e. in the host’s rest phase). Occasionally, the effect of misalignment depends on the pathogen’s own phase at inoculation – for example, starting infections with aligned trophozoites results in lower virulence compared to misaligned trophozoites in terms of RBC loss (O’Donnell *et al*, 2014). Whilst parasite phase (i.e. IDC stage) at the point of inoculation was standardised in our experiment, future research asking how phase of both parasite and host interact to modulate the effects of misalignment on disease severity in *Plasmodium* as well as other rhythmic pathogens, for e.g. African trypanosomes (Rijo-Ferreira *et al*, 2017), will be informative for knowing whether disrupting rhythms in either party is a useful intervention strategy.

That virulent parasites are less synchronous in aligned infections and do not experience an increase in synchrony when misaligned, fits with observations that human malaria (*P. falciparum*) infections, which cause severe pathology, are not always highly synchronous (Simpson *et al*, 2002; Dobaño *et al*, 2007; Touré-Ndouo *et al*, 2009). However, estimating synchrony in natural infections is very challenging; bias can be introduced by using stage proportions (Greischar *et al*, 2019). However, this bias overestimates synchrony in expanding infections, thus, because CW-VIR density increases faster than CW-0, we are likely to have overestimated synchrony of CW-VIR. This makes our finding that CW-VIR is less synchronous, a conservative result. If synchrony causes detrimental crowding (Greischar *et al*, 2014), this cost must be traded off against the benefit of timing for exploiting RBCs and/or nutrients that are rhythmic due to the host’s feeding-fasting rhythm (Prior *et al*, 2021). Parasites could minimise the cost of synchrony by increasing their preferred age range for host RBCs (for example by infecting reticulocytes as well as normocytes), thus expanding their pool of available host cells – and mathematical modelling suggests that virulent parasites do employ this strategy (Antia *et al*, 2008). Future studies could explore the optimal level of synchrony and how the costs of synchrony trade-off against the benefits of timing by comparing the consequences of desynchronisation (i.e. initiating infections with equal numbers of each IDC stage) and misalignment on genotypes that differ in virulence. Furthermore, synchrony between parasites could bring intrinsic benefits, for example, enhancing cell-cell communication when parasites are at the same IDC stage. Such benefits are likely small but could be tested for by conducting experiments in arrhythmic hosts without canonical circadian clocks which allow the intrinsic impacts of synchrony to be assessed.

Finding that misaligned infections shorten their IDC duration (Fig. 4C) adds to the body of evidence that *P. chabaudi* readjusts its rhythm to match the phase of host rhythms by speeding up development (Subudhi *et al*, 2020; O’Donnell *et al*, 2022). That this change was greater for CW-VIR – and brought its IDC schedule closer to that of aligned counterparts (Fig. 4D) – supports the hypothesis that virulent parasites are better able to recover from misalignment. This finding is in keeping with virulent genotypes being generally superior at tolerating within-host stressors including competition with co-infecting genotypes and antimalarial drugs (De Roode *et al*, 2005; Bell *et al*, 2006; Schneider *et al*, 2008, 2012). Why virulent parasites are better at rescheduling is unlikely to be due to positive density dependence because rescheduling rate is independent of density (O’Donnell *et al*, 2022). Perhaps virulent parasites can better harness plasticity to modulate the IDC rhythm, similar to their greater capacity to adjust traits such as the sex ratio of their gametocytes under challenging conditions (Reece *et al*, 2008). Plasticity in the IDC rhythm may have a genetic basis upon which selection acts, because both the duration (‘period’) and synchrony (‘amplitude’) of the IDC varies between *P. chabaudi* genotypes (Prior, 2017) and different strains of the human-infective malaria parasite (*P. falciparum*) vary in their period (Smith *et al*, 2020). IDC rhythms also become altered during the peak of infections when hosts are most symptomatic and their own rhythms become dysregulated (Prior, 2017; Prior *et al*, 2019; O’Donnell *et al*, 2022); perhaps an enhanced ability to cope with variation in host rhythms contributes to virulence.

## 5 Conclusion

Overall, we find modest support for the hypothesis that virulent parasites are less synchronous and better able to cope with the impacts of misalignment. The effects of misalignment on parasite density may be small because our sampling occurred over the full course of acute infections, yet parasites can realign the IDC rhythm within a few IDC cycles (O’Donnell *et al*, 2022). Tracking the prevalence of virulent parasite strains and identifying circumstances that favour them is of public health importance. Different IDC stages vary in their sensitivity to antimalarials (Cambie *et al*, 1991; Caillard *et al*, 1992; Skinner *et al*, 1996; Owolabi *et al*, 2021) and misalignment enhances artemisinin efficacy for trophozoites (Owolabi *et al*, 2021). Innovative interventions could manipulate parasite rhythms to mould them into their most tolerable form to hosts or to schedule the IDC for maximal efficacy of existing drugs. Our data here warn that virulent parasites could be doubly tenacious against such interventions as these parasites may be inherently less susceptible to conventional antimalarials (Schneider *et al*, 2008, 2012) and might bounce back faster from perturbations to their IDC rhythm than less virulent genotypes.

## Acknowledgements

We would like to thank Margaret Mackinnon for originally creating the parasite lines used in this study. We also thank members of the Reece lab (Alejandra Herbert Mainero, Jacob Holland, Ronnie Mooney, Aidan O’Donnell, Orsolya Sárkány and Mary Westwood) for help with data collection. This work was supported by the Wellcome Trust (108905/B/15/Z; 202769/Z/16/Z) and the Royal Society (UF110155).

## Data availability statement

The data *will be available upon publication* at the Edinburgh DataShare Repository: https://datashare.ed.ac.uk/.

## Ethics approval statement

All procedures occurred in accordance with the UK Home Office regulations (Animals Scientific Procedures Act 1986; SI 2012/3039; project licence number 70/8546) and were approved by The University of Edinburgh.

