## Supplementary material for "Virulence is associated with daily rhythms in the within-host replication of the malaria parasite *Plasmodium chabaudi*": Suppl.

### Supplementary Tables

**Supplementary Table 1. Host health before infection does not differ between treatment groups.**

| | *test statistic* | *p-value* |
|---|---|---|
| *(A) Baseline weight, day -1 PI* | | |
| Weight~Alignment×Genotype | | |
| Alignment×Genotype | $F_{(1,46)} = 0.58$ | $p = 0.449$ |
| Alignment | $F_{(1,47)} = 0.13$ | $p = 0.721$ |
| Genotype | $F_{(1,48)} = 2.06$ | $p = 0.158$ |
| *(B) Baseline RBC, day -1 PI* | | |
| RBC~Alignment×Genotype | | |
| Alignment×Genotype | $F_{(1,46)} = 0.08$ | $p = 0.775$ |
| Genotype | $F_{(1,47)} = 0.02$ | $p = 0.898$ |
| Alignment | $F_{(1,48)} = 2.45$ | $p = 0.141$ |

Full linear models and results for baseline weight and RBC density one day before infections. Alignment refers to whether the intraerythrocytic developmental cycle (IDC) is aligned or misaligned to host circadian rhythm, and Genotype refers to either the relatively avirulent CW-0 or the more virulent CW-VIR parasites. PI= post infection.

**Supplementary Table 2. Parasite fitness.**

| | | |
|---|---|---|
| *(A) Total parasite density dynamics days 3-16 PI* | *test statistic* | *p-value* |
| $\log_{10}$(Par dens)~Days PI×Alignment×Genotype+(1\|Mouse) | | |
| Days PI×Alignment×Genotype | $\chi^2$ = 16.73, df = 13 | *p* = 0.212 |
| Alignment×Genotype | $\chi^2$ = 0.33, df = 1 | *p* = 0.568 |
| **Days PI×Alignment** | **$\chi^2$ = 82.17, df = 13** | ***p* < 0.001** |
| **Days PI×Genotype** | **$\chi^2$ = 278.50, df = 13** | ***p* < 0.001** |
| *(B) Cumulative total parasite density days 3-16 PI* | | |
| $\log_{10}$(Cumulative Par dens)~Alignment×Genotype | | |
| Alignment×Genotype | $F_{(1,41)}$ = 0.51 | *p* = 0.480 |
| Alignment | $F_{(1,42)}$ = 0.62 | *p* = 0.437 |
| **Genotype** | **$F_{(1,43)}$ = 68.44** | ***p* < 0.001** |
| *(C) Gametocyte density dynamics days 3-16 PI* | | |
| $\log_{10}$(Gam dens)~Days PI×Alignment×Genotype+(1\|Mouse) | | |
| **Days PI×Alignment×Genotype** | **$\chi^2$ = 35.68, df = 13** | ***p* = 0.001** |
| *(D) Peak gametocyte density in first peak (days 3-8 PI)* | | |
| log10(Peak Gam dens)~Alignment×Genotype | | |
| Alignment×Genotype | $F_{(1,46)}$ = 1.07 | *p* = 0.307 |
| Genotype | $F_{(1,47)}$ = 2.08 | *p* = 0.156 |
| **Alignment** | **$F_{(1,48)}$ = 13.55** | ***p* < 0.001** |
| *(E) Peak gametocyte density in second peak (days 9-16 PI)* | | |
| log10(Peak Gam dens)~Alignment×Genotype | | |
| Alignment×Genotype | $F_{(1,41)}$ = 0.32 | *p* = 0.572 |
| Alignment | $F_{(1,42)}$ = 0.01 | *p* = 0.912 |
| **Genotype** | **$F_{(1,43)}$ = 22.55** | ***p* < 0.001** |
| *(F) Cumulative gametocyte density days 3-16 PI* | | |
| $\log_{10}$(Cumulative Gam dens)~Alignment×Genotype | | |
| Alignment×Genotype | $F_{(1,41)}$ = 1.03 | *p* = 0.317 |
| Alignment | $F_{(1,42)}$ = 4.01 | *p* = 0.052 |
| **Genotype** | **$F_{(1,43)}$ = 12.94** | ***p* < 0.001** |

Full models and results of linear mixed effects (A, C) and linear models (B, D, E, F) for total parasite and gametocyte densities, with significant terms included in the final models in bold. Alignment refers to whether the IDC is aligned or misaligned to host circadian rhythm, and Genotype refers to either the relatively avirulent CW-0 or the more virulent CW-VIR. PI= post infection. Note: For D and E, all mice contributed to the first gametocyte peak (n = 50) but mice which were euthanised during day 9-16 PI were excluded from the analysis of the second gametocyte peak (remaining n = 45).

**Supplementary Table 3. Disease severity.**

| | *test statistic* | *p-value* |
|---|---|---|
| *(A) Weight dynamics days 3-16 PI* | | |
| Weight~Days PI×Alignment×Genotype+(1\|Mouse) | | |
| Days PI×Alignment×Genotype | $\chi^2$ = 12.06, df = 13 | $p$ = 0.523 |
| Alignment×Genotype | $\chi^2$ = 0.28, df = 1 | $p$ = 0.599 |
| **Days PI×Alignment** | **$\chi^2$ = 55.07, df = 13** | **$p$ < 0.001** |
| **Days PI×Genotype** | **$\chi^2$ = 192.16, df = 13** | **$p$ < 0.001** |
| *(B) Cumulative weight days 3-16 PI* | | |
| Cumulative weight~Alignment×Genotype | | |
| Alignment×Genotype | $F_{(1,41)}$ = 0.62 | $p$ = 0.436 |
| Alignment | $F_{(1,42)}$ = 0.56 | $p$ = 0.458 |
| **Genotype** | **$F_{(1,43)}$ = 6.35** | **$p$ = 0.016** |
| *(C) RBC dynamics days 3-16 PI* | | |
| RBC~Days PI×Alignment×Genotype+(1\|Mouse) | | |
| Days PI×Alignment×Genotype | $\chi^2$ = 18.67, df = 13 | $p$ = 0.134 |
| Alignment×Genotype | $\chi^2$ = 0.04, df = 1 | $p$ = 0.850 |
| **Days PI×Alignment** | **$\chi^2$ = 120.68, df = 13** | **$p$ < 0.001** |
| **Days PI×Genotype** | **$\chi^2$ = 122.84, df = 13** | **$p$ < 0.001** |
| *(D) Cumulative RBC days 3-16 PI* | | |
| Cumulative RBC~Alignment×Genotype | | |
| Alignment×Genotype | $F_{(1,41)}$ = 0.16 | $p$ = 0.693 |
| **Alignment** | **$F_{(1,42)}$ = 11.94** | **$p$ = 0.001** |
| **Genotype** | **$F_{(1,42)}$ = 41.78** | **$p$ < 0.001** |

Full models and results of linear mixed effects (A, C) and linear models (B, D) for weight and anaemia, with significant terms remaining in the final models in bold. Alignment refers to whether the IDC is aligned or misaligned to host circadian rhythm, and Genotype refers to either the relatively avirulent CW-0 or the more virulent CW-VIR. PI= post infection.

**Supplementary Table 4. Intra-erythrocytic developmental cycle rhythmicity parameters.**

| | *test statistic* | *p-value* |
|---|---|---|
| *(A) Amplitude* | | |
| Amplitude~Alignment×Genotype | | |
| **Alignment×Genotype** | $\boldsymbol{F_{(1,14)}}$**= 38.64** | $\boldsymbol{p < 0.001}$ |
| *(B) Period* | | |
| Period~Alignment×Genotype | | |
| **Alignment×Genotype** | $\boldsymbol{F_{(1,14)}}$**= 13.55** | $\boldsymbol{p = 0.002}$ |
| *(C) Phase* | | |
| Phase~Alignment×Genotype | | |
| **Alignment×Genotype** | $\boldsymbol{F_{(1,14)}}$**= 10.06** | $\boldsymbol{p = 0.007}$ |

Full linear models and results for amplitude, period, and phase, with significant terms remaining in the final models in bold. Alignment refers to whether the IDC is aligned or misaligned to host circadian rhythm, and Genotype refers to either the relatively avirulent CW-0 or the more virulent CW-VIR.

### Supplementary Figures

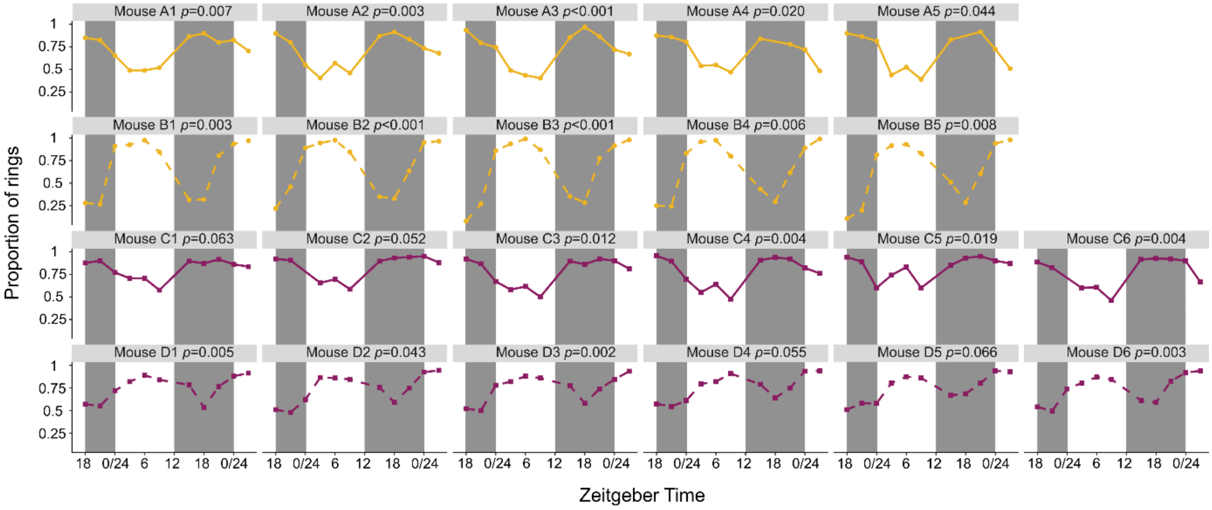


**Supplementary Fig. 1. Ring stage rhythm in individual mice.** Proportion of ring stage parasites in the blood from 42-75 hours PI for mice with aligned infections (solid lines) and 30-63 hours PI for mice with misaligned infections (dashed lines), for genotypes CW-0 (yellow) and CW-VIR (purple). Time is presented as Zeitgeber Time (ZT, i.e. time since light on). Grey shading from ZT12-24 indicates lights off (dark phase). All mice had rhythmic ring proportions according to the function *rain* ($p < 0.001$) and *p*-values for *meta2d* are indicated in the figure.

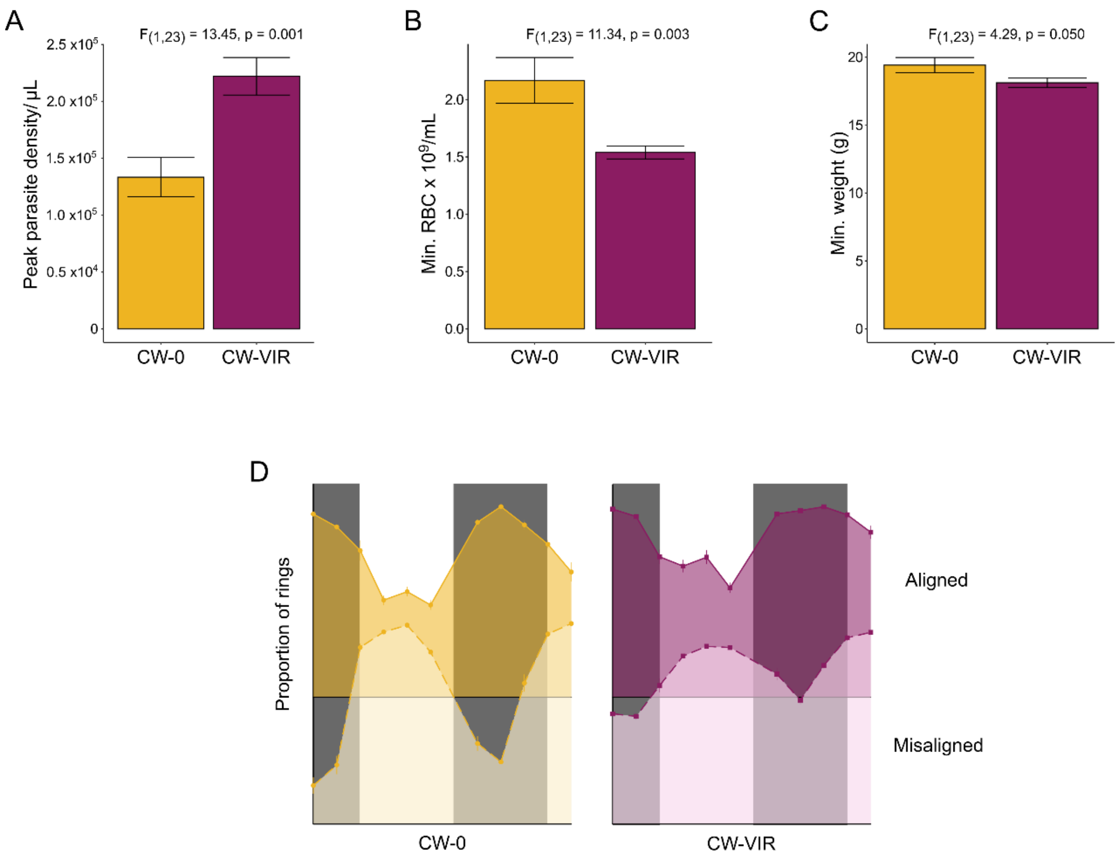


**Supplementary Fig. 2. The assumptions of the experimental design were met.** CW-VIR (N = 14, yellow) is more virulent than CW-0 (N=11, purple) in aligned infections, as shown by (**A**) higher peak parasite densities, (**B**) lower minimum red blood cell density and (**C**) borderline lower minimum weight for CW-VIR compared to CW-0. (**D**) Ring stages in misaligned groups (30-63 hours post infection, dashed lines) peak in opposite phases compared to aligned groups (42-75 hours post infection, solid lines), revealing that misalignment was successful. N = 5 CW-0 and N = 6 CW-VIR per alignment. Mean ± S.E.M are shown in all graphs. Grey shading from ZT12-24 indicates lights off (dark phase).
